# Image Reconstruction Reveals the Impact of Aging on Face Perception

**DOI:** 10.1101/2019.12.20.883215

**Authors:** Chi-Hsun Chang, Dan Nemrodov, Natalia Drobotenko, Adrian Nestor, Andy C. H. Lee

## Abstract

Extensive work has demonstrated a decline in face recognition abilities associated with healthy aging. To date, however, there has been limited insight into the nature and the extent of aging-related alterations in internal face representations. Here, we sought to address these issues by using an image reconstruction approach that capitalizes on the structure of behavioral data to reveal the pictorial content of visual representations. To this end, healthy young and older adults provided similarity judgments with pairs of face images. Facial shape and surface features were subsequently derived from the structure of the data for each participant and combined into image reconstructions of facial appearance. Our findings revealed that image reconstruction was successful for every participant, irrespective of age. However, reconstruction accuracies of shape and surface information were lower for older individuals than young individuals. Specifically, facial features diagnostic for face perception, such as eye shape and skin tone, were reconstructed poorly in older adults relative to young adults. At the same time, we found that age-related effects only accounted for a relatively small proportion of individual variability in face representations. Thus, our results provide novel insight into age-related changes in visual perception, they account for the decline in facial recognition occurring with age and they demonstrate the utility of image reconstruction to uncovering internal representations across a variety of populations.

---

Image Reconstruction Reveals the Impact of Aging on Face Perception Aging-related decrements in face recognition abilities have been extensively documented (Boutet, Taler, & Collin, 2015; Lee, Smith, Grady, Hoang, & Moscovitch, 2014; Rousselet et al., 2009; Thomas et al., 2008). Yet, the presumed alteration of internal face representations responsible for such effects is still unclear. In particular, the precise nature and the extent of the impact that aging has upon visual representations of facial identity remain to be elucidated.

A perception-based decline in face identification is apparent in healthy adults after 50 and especially after 70 years of age (Bowles et al., 2009). This decline is accompanied by complementary ones in recognition abilities including those in expression and social cue recognition (Halberstadt, Ruffman, Murray, Taumoepeau, & Ryan, 2011; Ruffman, Henry, Livingstone, & Phillips, 2008). Relevantly here, a decline in recognition performance is ascribed to decreasingly efficient visual representations, especially as related to holistic face processing (Chaby, Narme, & George, 2011; Wiese, Kachel, & Schweinberger, 2013), that is, the extent to which faces are encoded as an integrated whole, rather than a collection of different parts (Maurer, Le Grand, & Mondloch, 2002; Richler & Gauthier, 2014; Tanaka & Sengco, 1997). However, holistic-based accounts of this decline have been contested (Boutet & Faubert, 2006; Bowles et al., 2009). More importantly, the precise aspects of face representations undergoing deleterious age-related changes remain to be clarified.

Of particular relevance here is the distinction between facial shape and surface information (Bruce & Young, 1998), both of which are critical to face perception (Andrews, Baseler, Jenkins, Burton, & Young, 2016; Jiang, Blanz, & Toole, 2006; Jiang, Dricot, Blanz, Goebel, & Rossion, 2009; Michel, Rossion, Bülthoff, Hayward, & Vuong, 2013; Russell, Biederman, Nederhouser, & Sinha, 2007). Investigations of facial shape have targeted in particular second-order configural information, construed as the spatial layout of facial features (e.g., the distance between the eyes or between the nose and the mouth). For instance, several studies have artificially manipulated face shape, by modifying the layout of facial features (e.g., by changing the eye-to-mouth distance) and found impaired sensitivity to this type of shape information in older adults (Chaby, Narme, & George, 2011; Murray, Halberstadt, & Ruffman, 2010; Slessor, Riby, & Finnerty, 2013). However, this work has been typically constrained by the examination of a handful of predetermined features. Furthermore, it has yielded contradictory findings pointing to impaired holistic processing (Chaby et al., 2011), lack of such impairment (Boutet & Faubert, 2006) or increased holistic processing in older adults (Adduri & Marotta, 2009; Daniel & Bentin, 2012).

Regarding surface information, which includes color, texture and reflectance, even less is currently known. While decreasing contrast sensitivity in older adults has been found to adversely impact recognition (Barnes, De l’Aune, & Schuchard, 2011; Lott et al., 2005), it is unclear to what extent this is due to impaired surface versus shape processing. Thus, the contribution of shape information to the age-related decline in face identification is a topic of debate while the contribution of surface information is currently unknown.

Accordingly, the present work sought to provide novel insights into the issues above by employing image reconstruction methodology (Miyawaki et al., 2008; Nestor, Plaut, & Behrmann, 2016; Nishimoto et al., 2011; Thirion et al., 2006). Such methodology has been proved effective at retrieving the appearance of individual faces (Chang, Nemrodov, Lee, & Nestor, 2017; Nemrodov, Behrmann, Niemeier, Drobotenko, & Nestor, 2019; Zhan, Garrod, van Rijsbergen, & Schyns, 2019) from perception and memory with the aid of behavioral and neuroimaging data. Briefly, this approach capitalizes on the structure of face space (Valentine, 1991), a multidimensional construct which encodes individual faces as points and their perceptual similarity as the pairwise distance of corresponding points. In the present study, face space constructs were estimated separately for each participant from behavioral estimates of pairwise face similarity. Then, facial shape and surface features were derived from the structure of face space (Nemrodov et al., 2019) and combined to recreate the overall appearance of individual faces as perceived by young and older adults.

Key strengths of our approach include the data-driven aspect of image reconstruction (i.e., as opposed to the testing of a few facial features driven by specific hypotheses), the level of granularity with which visual content can be extracted (i.e., locally and across different visual attributes) as well as, critically, the ability to provide a conceptual and methodological framework for assessing the role of aging in face perception. Thus, the present study represents a significant advance by shedding light on age-related changes to face representations, their nature, their extent and their fine-grained content.

## Method

### Participants

Twenty-five Caucasian young adults (YAs) and 35 Caucasian older adults (OAs) participated in the study in exchange for payment or course credit. In total, 11 OAs were excluded, of which nine scored less than 26/30 on the Montreal Cognitive Assessment (MoCA; Nasreddine et al., 2005) and were subsequently considered at risk for mild cognitive impairment (MCI), one did not follow the correct experimental protocol, and one did not complete the whole study. Also, four YAs were excluded, of which three did not follow correctly the experimental instructions, and one did not complete the whole study. The final sample of participants included in the analyses below consisted of 21 YAs (11 female; *M*_age_ = 21.90 years; *SD*_age_ = 3.27; *M*_education_ = 15.57 years; *SD*_education_ = 1.96) and 24 OAs (11 female; *M*_age_ = 66.46 years; *SD*_age_ = 2.67; *M*_education_ = 16.46 years; *SD*_education_ = 3.31). YAs and OAs were significantly different in age (*t*(43) = −50.17, *p* < .001, Cohen’s *d* = 15.30, 95% CI of the difference [−46.34, −42.76]) but not education (*t*(43) = −1.07, *p* = .29). This sample size was determined on the basis of prior work on aging and face perception research (Lee et al., 2014; Norton et al., 2009).

All participants had normal or corrected-to-normal vision and no history of neurological or visual disorders. Informed written consent was obtained from all participants. All procedures were carried out in accordance with University of Toronto Research Ethics Guidelines and were approved by the University of Toronto Research Ethics Board.

### Experimental Procedures

#### Overview

Both groups of participants completed two experimental sessions, each lasting 2 to 2.5-hours, on different days separated at most by one week. Participants were administered a battery of tests including a series of standardized and experimental neuropsychological tests, followed by the main experiment.

All computerized neuropsychological tests as well as the main experimental task relied on Matlab (Mathworks, Natick, MA) and Psychtoolbox 3.0.12 (Brainard, 1997; Kleiner, Brainard, & Pelli, 2007; Pelli, 1997). Experimental instructions and stimuli were presented on a touch-sensitive LCD screen at a resolution of 1280 × 720 pixels.

#### Neuropsychological tests

Participants completed a series of tests including the MoCA (Nasreddine et al., 2005), the Logical Memory subtest from the Wechsler Memory Scale (WMS; 4th ed.; Wechsler, 2009), the Vocabulary and Similarity subtests from the Wechsler Abbreviated Scale of Intelligence (WASI; 2nd ed.; Wechsler, 2011), the Category Comprehension Test (word-picture matching from the semantic memory battery in Hodges & Patterson, 1995), the Short Recognition Memory Test for Faces and Words and the Topographical Recognition Memory Test from the Camden Memory Test (CMT; Warrington, 1996), the Copy and the Immediate Recall subtests from the Rey Complex Figure Test (RCFT; Meyers & Meyers, 1995), the Cambridge Face Memory Test (CFMT; Duchaine & Nakayama, 2006), as well as the face oddity judgment task (Lee et al., 2005) detailed below.

The face oddity judgment task (FOJT) is a computerized task that requires participants to distinguish the identities of multiple face stimuli by selecting the “odd” face from an array of faces. Previous work (Behrmann, Lee, Geskin, Graham, & Barense, 2016; Lee et al., 2005, 2006) has found that performance on this perceptual task is compromised in individuals with medial temporal lobe damage as well as in individuals with congenital prosopagnosia.

In detail, each trial on this test started with a central fixation cross (500 ms), followed by an array of four face images presented against a grey background - three of the same identity and one of a different identity. Participants were instructed to select the “odd” face by touching it on the screen and face images remained visible until participants made a response. Stimuli consisted of grayscale images of 80 facial identities (all male aged 20-40 years, with short hair, no facial hair or spectacles). Each facial identity was captured from 4 different views (looking directly ahead, 45° to the right, 45° to the left, and upwards). Two versions of the task were administered: in the same-view task all four faces in a trial were presented from the same viewpoint whereas in the different-view task, each face was presented from a different viewpoint. Each version of the task consisted of 40 trials, with no stimulus repetition, preceded by 4 practice trials (using 2 separate facial identities).

#### Main experimental task

The main experiment relied on a pairwise similarity rating task with faces of different identities. Each trial started with a central fixation cross (500 ms), followed by a pair of face images presented side by side against a dark background. Each face subtended an angle of 2.7° x 4° from 97 cm and was displaced 2.3° from the center of the screen. Participants were asked to rate the similarity of the two faces on a 7-point scale and to verbally report their rating. The experimenter then recorded their response by pressing a corresponding number key, which led to the end of the trial. The left/right location of the images was counterbalanced and each face was paired with every other face exactly once, leading to 1596 trials presented in random order and divided equally over 14 blocks. Stimuli consisted of 57 color face images selected from four databases, including Radbound (Langner et al., 2010), AR (Martinez & Benavente, 1998), FEI (Thomaz & Giraldi, 2010) and FERET (Phillips, Moon, Rizvi, & Rauss, 2000; Phillips, Wechsler, Huang, & Rauss, 1998). All of them displayed unfamiliar faces of Caucasian males with a neutral expression from a frontal viewpoint and with frontal lighting. In addition, face images were cropped to reveal only the internal features of the face, spatially normalized with the position of the eyes and the nose and color-normalized with the same mean and contrast values separately for each CIEL*a*b color channel. Twelve practice trials were administered at the start of the task.

### Analyses

#### Neuropsychological test analyses

For each standardized test, the raw scores for YAs and OAs were compared using an independent-sample t-test or a permutation test when data were not normally distributed (Shapiro-Wilk test, *p* < .05 for MoCA, Category Comprehension, CMT topography, CMT face, CMT word, RCFT copy, same-view FOJT). In the permutation test, group labels were randomly shuffled 1,000 times and the difference of the means between two groups was computed for each permutation, resulting in a distribution of mean differences. A significance value for the true difference was then obtained by computing the proportion of permutation-based differences larger than or equal to it.

For FOJT, accuracy (proportion correct) was first analyzed using a two-way mixed-design analysis of variance (ANOVA), with a between-subject factor of age (YA vs. OA) and a within-subject factor of viewpoint (same vs. different). Significant interactions were further explored by pairwise group comparisons (i.e., independent-sample t-test or permutation test as described above for non-normally distributed data (Shapiro-Wilk test, *p* < .05)).

#### Image reconstruction

The procedure follows a recent approach to facial image reconstruction (Nemrodov et al., 2019) which relies on face de/re-composition into shape and surface information. Specifically, the procedure separately recovers shape, defined as the position of a set of fiducial points corresponding to key facial features (i.e., 82 points x 2 in-plane coordinates), and surface, as given by pixel intensity values in CIEL*a*b* after warping face images to a standard shape template.

Briefly, the procedure involves a sequence of steps, as follows: (a) each face stimulus was decomposed into shape and surface information with the aid of the InterFace toolbox (Kramer, Jenkins, & Burton, 2017); (b) a 20-dimensional face space construct (Figure 1c) was estimated by applying multidimensional scaling (MDS) to a confusability matrix based on pairwise similarity of faces obtained from similarity ratings (Figure 1a), excluding the target face (image **n** in Figure 1b); (c) classification vectors (CVs), for shape, and classification images (CIMs), for surface (Figure 1d), were synthesized through a method akin to reverse correlation (Murray, 2011; Neri & Levi, 2006; Smith, Gosselin, & Schyns, 2012) from all faces but the target by computing a weighted average of face images proportionally with their coordinates in MDS-based space; (d) the significance of each CV and CIM was assessed via a pointwise/pixelwise permutation test (i.e., by randomizing images with respect to their coordinates on each dimension and by recomputing CVs or CIMs for a total of 1,000 shuffles; two-tailed t-test; FDR correction: *q* < 0.1); (e) the target face (Figure 1c) was projected in the existing face space based on its perceived similarity to the other faces; (f) the shape and surface of the target face were recovered through a linear combination of significant CVs or CIMs, proportionally to the coordinates of the target in face space, added onto an average shape and surface (Figure 1d); and, last, (g) the recovered shape and surface were combined into a reconstructed face using the InterFace toolbox (Kramer et al., 2017) (Figure 1d).

**Figure 1.**
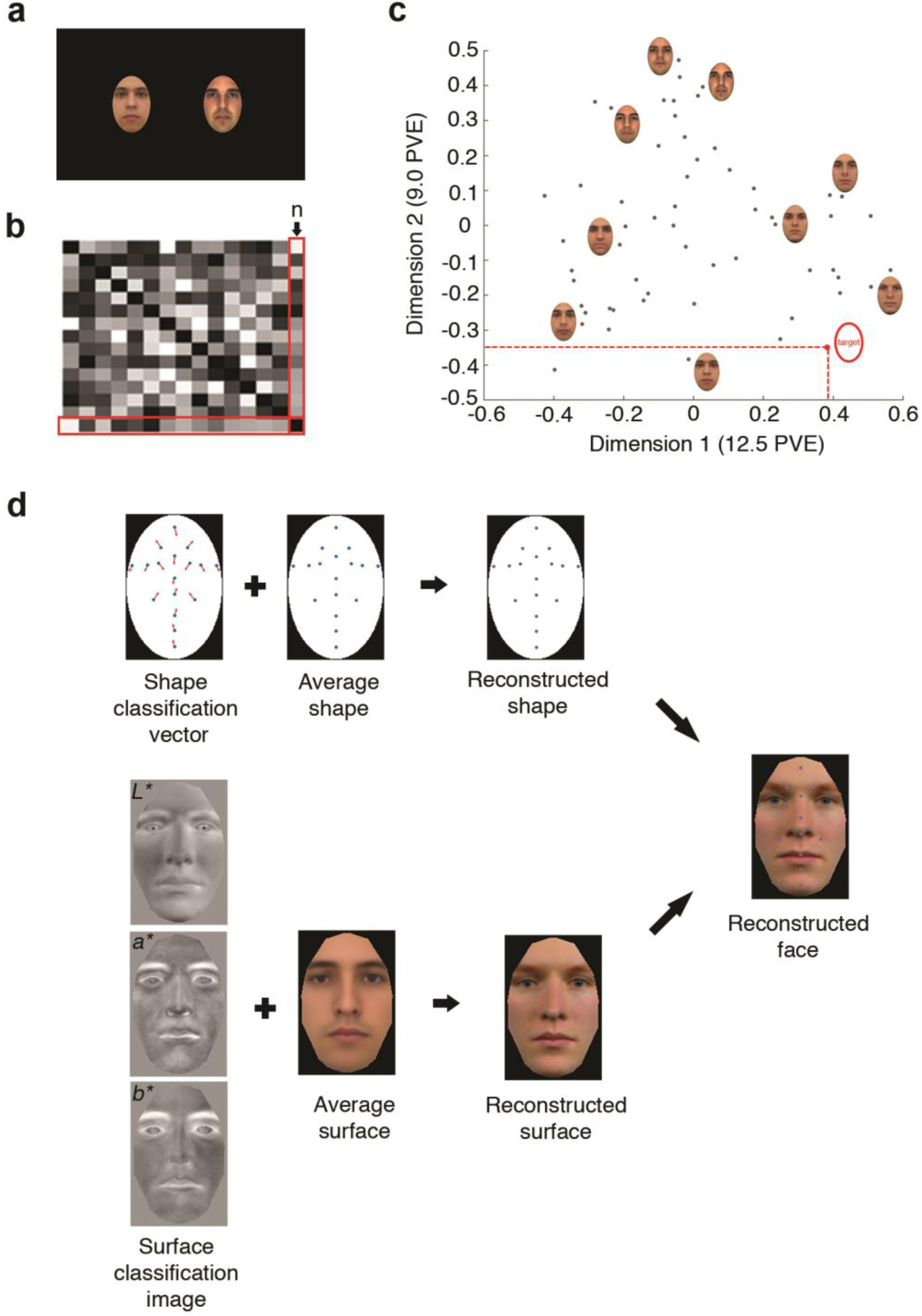
Reconstruction procedure: (a) participants rate the similarity of pairs of faces that are simultaneously presented on the screen; (b) pairwise similarity ratings are converted into a confusability matrix consisting of **n** faces, where face **n** is the target face for reconstruction; (c) a 20-dimensional face space is constructed by applying the multidimensional scaling to the similarity of **n-1** faces in the confusability matrix and the coordinates of the target face **n** in the face space is estimated (only the first 2 dimensions are presented for illustration purpose; PVE – percentage of variance explained); (d) classification vectors (CVs) for shape, or classification images (CIMs) for surface are derived via image classification and analyzed using a pointwise or pixelwise permutation test (FDR-corrected; *q* < 0.1); subsequently, shape and surface properties of the target face are separately reconstructed through a combination of significant CVs or CIMs added onto an average shape and surface; last, shape and surface are recombined into a reconstructed face. (Due to copyright restrictions, all images of original face stimuli have been replaced with face images artificially generated in Matlab; thus, they do not display photographs of actual persons. The reconstructed surface and reconstructed face in (**d**) are generated by our image reconstruction algorithm in Matlab and InterFace.)

It important to highlight that the leave-one-out procedure above enforces non-circularity by excluding the reconstruction target from the estimation of face space and of its underlying features. Also, we note that facial features are synthesized directly from the structure of the experimental data, rather than pre-selected based on prior hypotheses.

#### Evaluation of reconstruction results

Reconstruction accuracy was measured based on the pointwise similarity, for shape, or pixelwise similarity, for surface, between reconstructed images and target faces. Specifically, each reconstructed shape was compared to its target via the Euclidean distance computed across all corresponding 82 fiducial point coordinates. Then, its reconstruction accuracy was measured as the proportion of instances for which that distance was smaller than the distance between reconstruction and the stimulus shapes of all other stimuli but the target. Surface accuracy was measured similarly, except that Euclidean distances were computed across pixel values in CIEL*a*b* space. The latter procedure was also followed to measure the accuracy of reconstructed faces.

Next, for each type of reconstruction (i.e., shape, surface or reconstructed faces) accuracies were averaged across items and success was assessed relative to chance (50%) using a permutation test, separately for each participant. To this end, for each target face, the positions of the other faces in MDS-based space were randomly shuffled 1,000 times and the reconstruction of the target was recomputed. Then, significance was computed as the proportion of instances for which true reconstruction accuracy was higher than or equal to its permutation-based counterparts.

Further, to examine group-level effects, reconstruction was evaluated using a two-way mixed-design ANOVA, with one between-subject factor of age (YA vs. OA) and one within-subject factor of facial attribute (shape vs. surface). Also, reconstructed faces of YAs and OAs were compared using an independent-sample t-test.

To examine in more detail the impact of aging on color processing, average surface reconstruction accuracy was computed separately for each of three color channels in the CIEL*a*b color space, where L*, a* and b* represent the luminance, red-green, and yellow-blue color-opponent channels of human vision. Accuracy was then assessed using a two-way mixed-design ANOVA, with one between-subject factor of age (YA vs. OA) and one within-subject factor of color channel (L*, a*, and b*).

Next, to elucidate whether aging impacts the perception of specific facial features, heatmaps highlighting the differences in reconstruction accuracy between YAs and OAs were constructed and examined separately for shape and surface. Specifically, we calculated, for each participant the reconstruction accuracy of each fiducial point, for shape, and of each pixel intensity value, for surface. Then, the reconstruction accuracy of each fiducial point and pixel intensity value was averaged across both sides of a stimulus to boost the robustness of the estimates and the significance of the difference between groups was evaluated via a fiducial-pointwise or pixelwise parametric test (two-tailed t-test; FDR correction: *q* < 0.05).

Last, reconstruction accuracies were correlated with the scores of neuropsychological tests associated with face processing, including the Short Recognition Memory Test for Faces from CMT, CFMT, and the FOJT, allowing us to examine if and how reconstruction results are related to more general recognition abilities across participants.

#### Aging and individual differences

To investigate the nature of individual variability in face representations, including but not limited to the role of aging, we conducted a sequence of analyses as follows.

First, we applied principal component analysis (PCA) to the pairwise confusability scores among all faces across all YA and OA participants. This allows us to construct a participant space reflecting individual differences in how similarly different faces are perceived. Specifically, each participant was represented as a point in a 10-dimensional PCA-based space. This number of dimensions was selected under the twofold constraint of keeping dimensionality low, to prevent the overfitting of classification models in this space, and to account for at least half of data variance.

Second, to assess how robustly age serves to segregate participants in this space, we classified participants based on age group with the aid of linear discriminant analysis (LDA) and a leave-one-out cross-validation schema. Specifically, all participants, but one, were used to train an LDA classifier and the resulting model was tested on the left-out participant (i.e., represented as a 10-dimensional vector). Classification accuracy was tested by systematically leaving out each participant and significance was established with the aid of permutation tests (based on 1,000 shuffles of participant location in PCA space).

Third, heatmap computation for PC1 and PC2 was conducted separately for shape and surface to reveal facial features that are associated with individual difference in face perception. These heatmaps were constructed by using the same procedure for constructing the heatmaps of difference in reconstruction accuracy between age groups, with the exception that participants were grouped by the PC scores (i.e., positive vs. negative) rather than age.

## Results

### Neuropsychological Tests

Standardized tests revealed significantly better performance for YAs relative to OAs across several tests: WASI-II similarity, RCFT copy and immediate recall as well as CFMT (see Table 1 for statistics). For FOJT, a two-way mixed-design ANOVA showed a main effect of age (*F*(1,43) = 4.91, *p* = .03, *η*^2^ = .10), viewpoint (*F*(1,43) = 128.18, *p* < .001, *η*^2^ = .70), and an interaction between the two factors (*F*(1,43) = 11.85, *p* = .001, *η*^2^ = .06). Specifically, YAs performed significantly better than OAs in the different-view condition, but significantly worse in the same-view condition (see Table 1). However, we note that performance of both groups in the same view condition was close to ceiling (> 97% accuracy) and the significant difference between groups here was driven by several YA participants making a few errors whereas most OA participants attained a perfect score. In contrast, in the different-view condition the performance for both groups was not at ceiling and the difference between groups was considerably larger.

**Table 1.** Summary of Neuropsychological Tests Results of Young Adult and Older Adult Groups

| Test | Age groups |  | Statistics |
| --- | --- | --- | --- |
|  | YA | OA |  |
| MoCA (/30) | 28.00 (1.14) | 27.63 (1.25) | $D = 0.37, p = .35,$<br>95% CI = [-0.70, 0.65] |
| WMS-IV LM immediate recall (/50) | 24.90 (5.51)<br>50%ile | 26.75 (5.15)<br>63%ile | $t(43) = -1.16, p = .252,$<br>Cohen's $d = 0.35,$<br>95% CI = [-5.05, 1.36] |
| WMS-IV LM delayed recall (/50) | 21.71 (6.17)<br>50%ile | 23.04 (4.60)<br>63%ile | $t(43) = -0.82, p = .414,$<br>Cohen's $d = 0.25,$<br>95% CI = [-4.58, 1.92] |
| WMS-IV LM recognition (/30) | 24.62 (2.56)<br>50%ile | 24.29 (3.11)<br>50%ile | $t(43) = 0.38, p = .704,$<br>Cohen's $d = 0.11,$<br>95% CI = [-1.40, 2.06] |
| WASI-II vocabulary (/59) | 41.14 (5.98)<br>69%ile | 42.50 (4.98)<br>73%ile | $t(43) = -0.83, p = .411,$<br>Cohen's $d = 0.25,$<br>95% CI = [-4.65, 1.94] |
| WASI-II similarity (/45) | 36.00 (2.51)<br>79%ile | 33.42 (3.04)<br>69%ile | $t(38) = 2.94, p = .006,$<br>Cohen's $d = 0.93,$<br>95% CI = [0.80, 4.36] |
| Category comprehension (/64) | 63.33 (0.86) | 63.75 (0.68) | $D = -0.42, p = .09,$<br>95% CI = [-0.42, 0.48] |
| CMT topography (/30) | 25.57 (2.50)<br>50%ile | 26.08 (3.02)<br>75%ile | $D = -0.51, p = .54,$<br>95% CI = [-1.58, 1.54] |
| CMT face (/25) | 24.43 (0.98)<br>75%ile | 23.71 (1.55)<br>75%ile | $D = 0.72, p = .07,$<br>95% CI = [-0.72, 0.81] |
| CMT word (/25) | 24.95 (0.22)<br>90%ile | 24.83 (0.38)<br>90%ile | $D = 0.12, p = .36,$<br>95% CI = [-0.15, 0.21] |
| RCFT copy (/36) | 33.67 (2.13) | 30.61 (5.09) | $D = 3.06, p = .01,$ |
|  | > 16%ile | > 16%ile | 95% CI = [-2.18, 2.33] |
| RCFT immediate recall (/36) | 23.52 (4.33) | 12.78 (7.37) | $t(42) = 5.82, p < .001$, |
| | 38%ile | 34%ile | Cohen's $d = 1.76$, |
|  |  |  | 95% CI = [7.02, 14.47] |
| CFMT (/100%) | 79.30% | 68.06% | $t(43) = 3.33, p = .002$, |
| | (10.62%) | (11.83%) | Cohen's $d = 1.00$, |
|  | 58%ile | 43%ile | 95% CI = [0.04, 0.18] |
| Same-view FOJT (/100%) | 97.50% (2.37%) | 98.96% (1.94%) | $D = -0.015, p = .03$, |
|  |  |  | 95% CI [-0.012, 0.01] |
| Different-view FOJT (/100%) | 85.71% (9.72%) | 76.87% | $t(43) = 2.85, p = .007$, |
| | | (10.92%) | Cohen's $d = 0.85$, |
|  |  |  | 95% CI [0.03, 0.15] |

*Note.* Proportion correct responses are reported for Cambridge Face Memory Test (CFMT) and face oddity judgment task (FOJT), while average raw scores are reported for other neuropsychological tests, separately for young adult (YA) and older adult (OA) groups. Standard deviations are reported in the parentheses and percentile (%ile) scores, relative to the corresponding group norms, were also given, where available. Results shown in the third column are based on independent-sample t-tests, or from permutation tests when data is not normally distributed. MoCA = Montreal Cognitive Assessment; WMS-IV LM = Wechsler Memory Scale, 4th ed. Logical Memory subtest; WASI-II = Wechsler Abbreviated Scale of Intelligence, 2nd ed.; CMT = Camden Memory Test; RCFT = Rey Complex Figure Test; CI = confidence interval of the difference between age groups; D = value of the difference between age groups.

### Facial Image Reconstruction

An examination of single participant results revealed that reconstruction accuracy was significantly above chance for every YA and OA participant (permutation tests; all *p*s < .001). Examples of reconstructed images, relying on the combination of reconstructed shape and surface, are presented in Figure 2.

**Figure 2.**
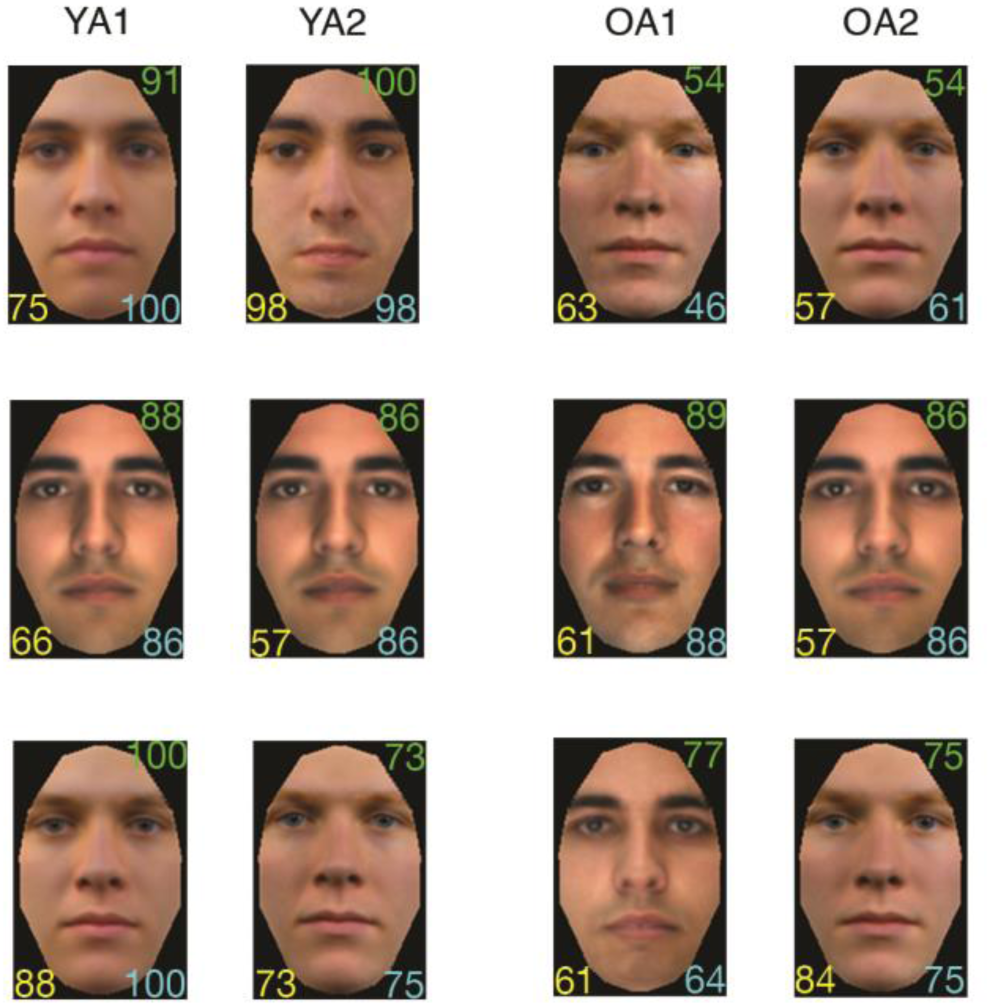
Examples of reconstructions (reconstructed faces) for two representative participants from each age group (YA – young adult; OA – older adult). Reconstruction accuracy is calculated separately for the reconstructed face (top right), the reconstructed shape (bottom left), and the reconstructed surface (bottom right). (Corresponding stimuli images for the reconstructions could not be reproduced due to copyright restrictions; all reconstructed faces were generated by combining reconstructed shape and reconstructed surface in InterFace)

A two-way ANOVA (between-subject age factor: YA and OA x within-subject attribute factor: shape and surface) revealed a main effect of age (*F*(1,43) = 12.35, *p* = .001, *η*^2^ = .23) (Figure 3a), with reconstruction accuracies being greater for YA compared to OA for both shape and surface (*M*_YA_ = 0.69; *M*_OA_ = 0.66). The main effect of facial attribute was also significant (*F*(1,43) = 1004.93, *p* < .001, *η*^2^ = .96), with surface information being more accurately retrieved than shape information for both groups of participants (*M*_shape_ = 0.61; *M*_surface_ = 0.74). No effect of interaction between age and facial property was found (*F*(1,43) = 0.094, *p* = .76).

**Figure 3.**
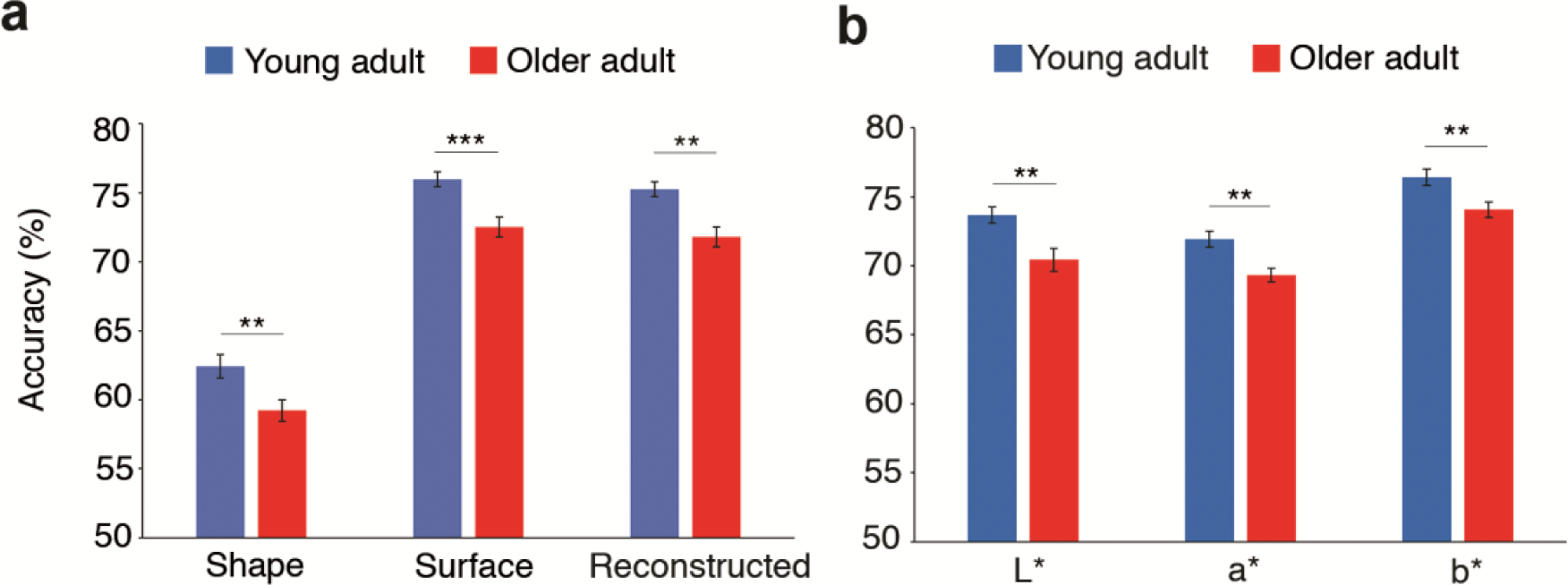
Average reconstruction accuracy for each age group. Accuracy is estimated for (a) shape, surface and reconstructed faces, as well as for (b) each CIEL*a*b* color channel (error bars ±1 *SE*; \*\**p* < .01, \*\*\**p* < .001)

A two-way ANOVA applied to reconstruction accuracy for each color channel (between-subject age factor: YA and OA x within-subject color factor: L*, a* and b*) found a main effect of age (*F*(1,43) = 11.94, *p* = .001, *η*^2^ = .22) (Figure 3b), with the accuracy of every color channel higher for YA than OA (*M*_YA_ = 0.74; *M*_OA_ = 0.71). A main effect of color channel was also present (*F*(2,86) = 101.52, *p* < .001, *η*^2^ = .70), and further pairwise comparisons revealed that yellow-blue color information achieved the highest accuracy compared to the other two channels (b* vs. L*: *t*(44) = 9.86, *p* < .001, Cohen’s *d* = 1.47, 95% CI [0.02, 0.04]; b* vs. a*: *t*(44) = 16.54, *p* < .001, Cohen’s *d* = 2.47, 95% CI [0.04, 0.05]; Holm-Bonferroni-corrected), while luminance information was recovered more accurately than red-green information (*t*(44) = 3.72, *p* < .001, Cohen’s *d* = 0.55, 95% CI [0.01, 0.02], Holm-Bonferroni-corrected). No interaction effect was found between age and color channel (*F*(2,86) = 0.95, *p* = .39).

Accuracy heatmaps of reconstruction differences between YAs and OAs revealed that shape information was better retrieved in YAs especially for the eyes, but also for the nose (Figure 4a). Regarding surface information, differences between YAs and OAs were found in all three color channels (Figure 4b). Consistent with our analyses above (Figure 3b), differences were more widely spread for the yellow-blue channel relative to the other channels. In particular, the color of the forehead and cheeks was better retrieved for YAs than OAs. Thus, reconstruction results suggest that both shape and surface information processing are impacted by aging. Further, specific facial features diagnostic of facial identity, such as eye shape, were compromised in OAs.

**Figure 4.**
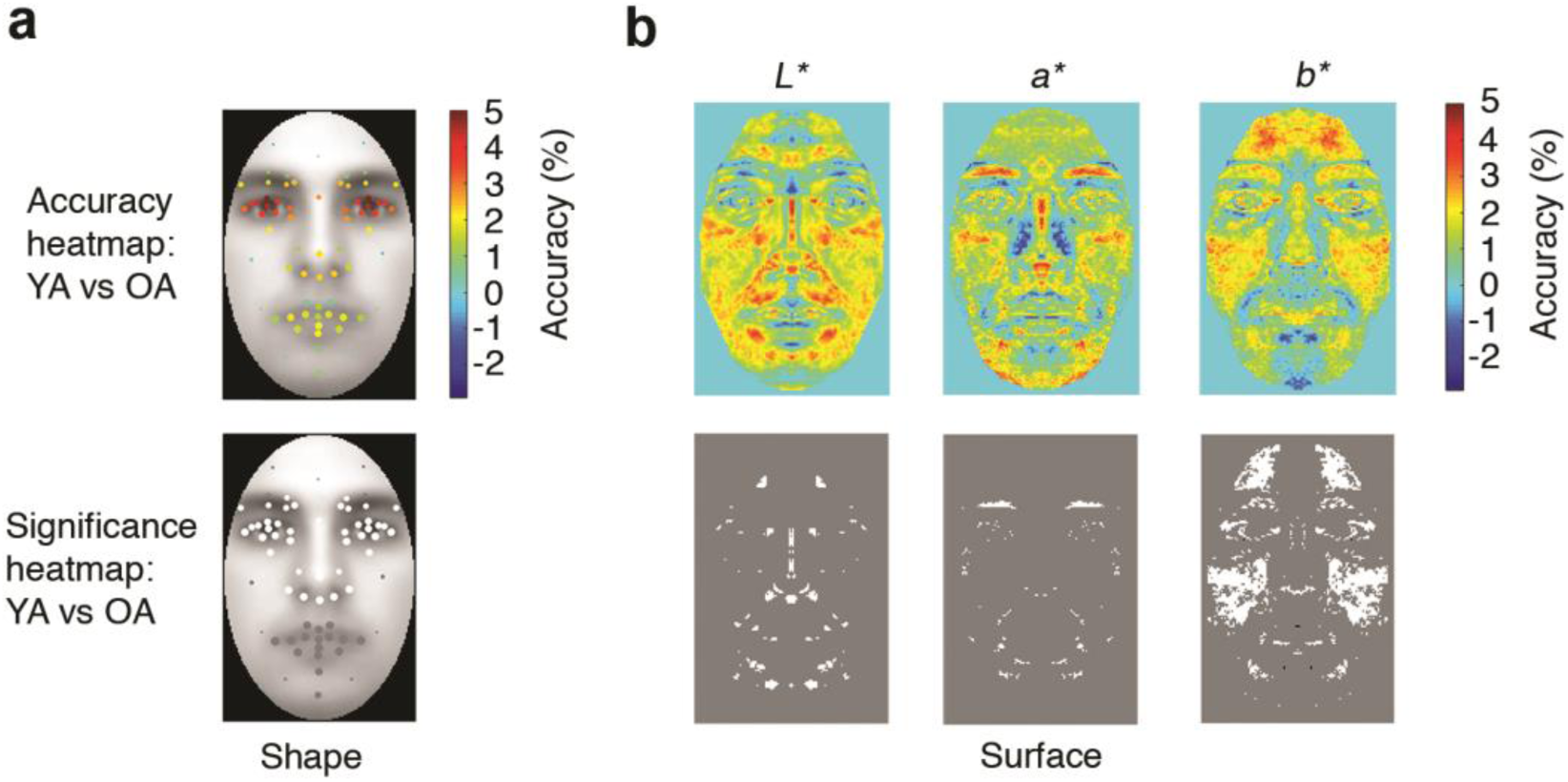
Differences in reconstruction accuracy between the two age groups (YA – young adult; OA – older adult). Heatmaps of such differences and their significance are constructed for (a) every fiducial point in shape, and (b) for every pixel in each color channel. In significance heatmaps, white indicates information more accurately retrieved in YA while black indicates information more accurately retrieved in OA (FDR, *q* < .05) across fiducial points for shape and pixels for surface (gray marks the lack of significant differences). The relative size of fiducial points corresponds to their location variability across faces (*SD*). (The gray face images in (**a**) were artificially generated for illustration purposes in Matlab)

Next, correlations between the scores of neuropsychological tests and reconstruction accuracy across all participants from both groups showed significant relationships between the accuracies of different-view FOJT and shape reconstruction (*r*(43) = .42, *p* = .03, Holm-Bonferroni-corrected) (Figure 5a), as well as surface reconstruction (*r*(43) = .48, *p* = .006, Holm-Bonferroni-corrected) (Figure 5b). Notably, the results remained significant even after controlling for age (different-view FOJT accuracy and shape reconstruction: *r*(42) = .31, *p* = .04; different-view FOJT accuracy and surface reconstruction: *r*(42) = .35, *p* = .04; both Holm-Bonferroni-corrected).

**Figure 5.**
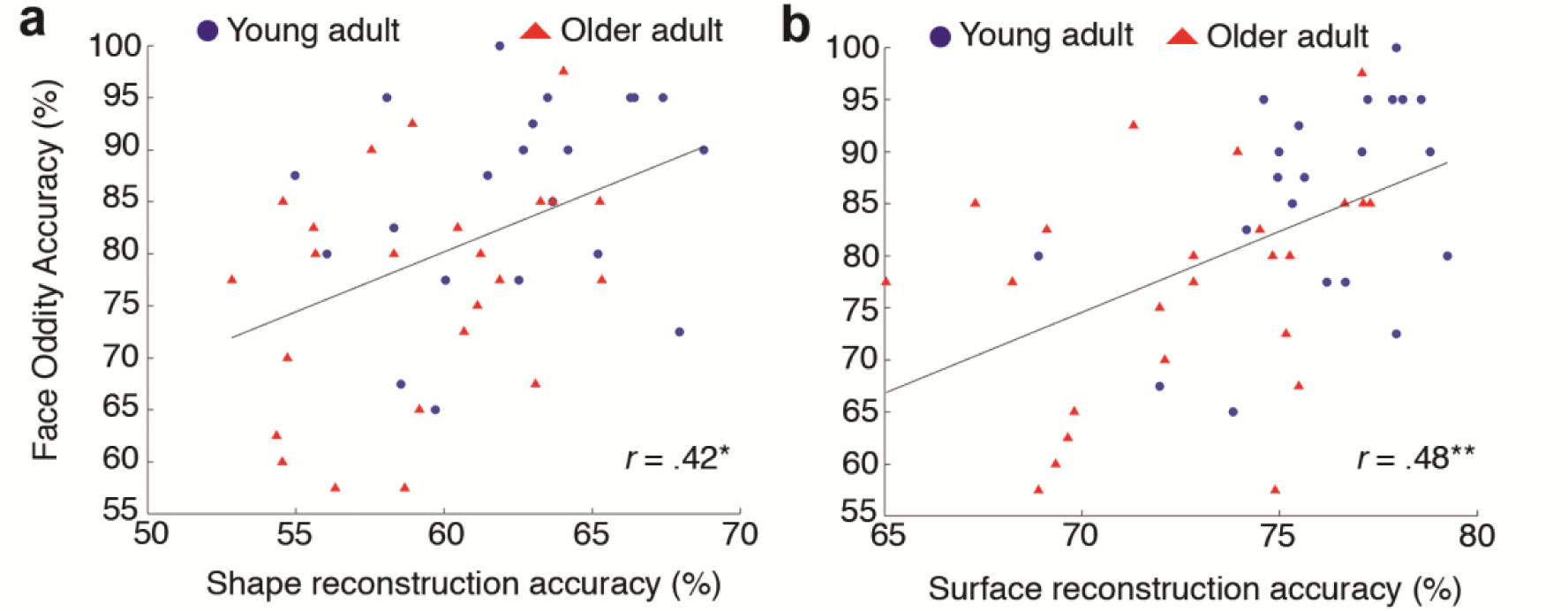
The relationship between different-view face oddity accuracy and (a) shape reconstruction accuracy as well as (b) surface reconstruction accuracy across the two age groups. Black lines indicate fitted linear trends (* *p* < .05; \*\**p* < .01).

Last, the results of reconstructed faces revealed higher reconstruction accuracy for YA than OA (*t*(43) = 3.68, *p* = .001, Cohen’s *d* = 1.12, 95% CI of the difference [0.02, 0.05]) (Figure 3a). This is consistent with the finding that both shape and surface information was retrieved better for YAs than OAs.

### Individual Variability and Aging

A multidimensional participant space was generated from the application of PCA to individual ratings of face similarity – the space was constrained to the first 10 dimensions, accounting for 51.29% data variance, out of which the first three are shown in Figure 6. An examination of this space revealed a clear segregation of participants based on age (especially along dimension 3 in Figure 6).

**Figure 6.**
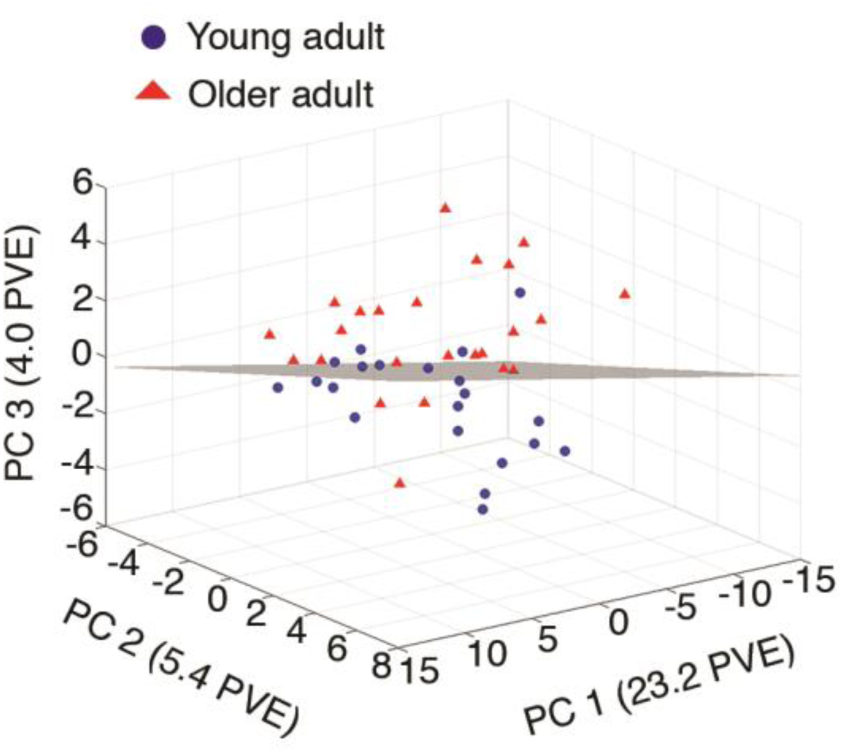
Multidimensional participant space. Each point represents a participant in the study. The space was derived from principle component analysis (PCA) as applied to pairwise confusability scores. The grey shaded plane is the hyperplane created by a Linear Discriminant Analysis classifier that distinguishes participants from different age groups. For convenience, only the first three dimensions are shown here (PC – principle component; PVE – percentage of variance explained).

To confirm and to quantify this segregation, LDA classification of age groups was conducted across participants using a leave-one-subject-out cross-validation procedure. This analysis found that participant age could be classified well above chance: 85.55% accuracy (permutation test; *p* < .001, 95% CI [35.45%, 66.65%]). This indicates that age has a significant impact on the structure of behavioral data, converging with the observed differences in image reconstruction between YAs and OAs, and serves to quantify this impact.

To further clarify the factors responsible for the structure of the space above, PC scores were correlated across participants with age as well as with performance accuracy based on neuropsychological tests. An examination of the first three dimensions displayed in Figure 6 found that age was significantly correlated only with scores on the third dimension (*r*(43) = .48, *p* = .01, Holm-Bonferroni-corrected), whereas accuracy in the different-view FOJT was correlated with performance on the first and second dimension scores (*r*(43) = .34, *p* = .02 and *r*(43) = .40, *p* = .007, uncorrected - see Figure S1 in the supplemental materials). Note that although the correlations between the first two PC scores and the different-view FOJT were not significant after correction for multiple comparisons (*p* = .28 and *p* = .10 for the PC1 and PC2, respectively), their effect size were between medium and large (Cohen, 1992).

Additionally, heatmaps of reconstruction accuracy for the first two PCs were constructed to investigate specific facial features that account for the structure underlying the participant space. The results showed that the PC1 captures surface information in all three color channels but not shape information (Figure 7a, b). In contrast, PC2 encodes primarily information about the shape of eyes and nose as well as red-green and yellow-blue color information, but no luminance information (Figure 7c, d).

**Figure 7.**
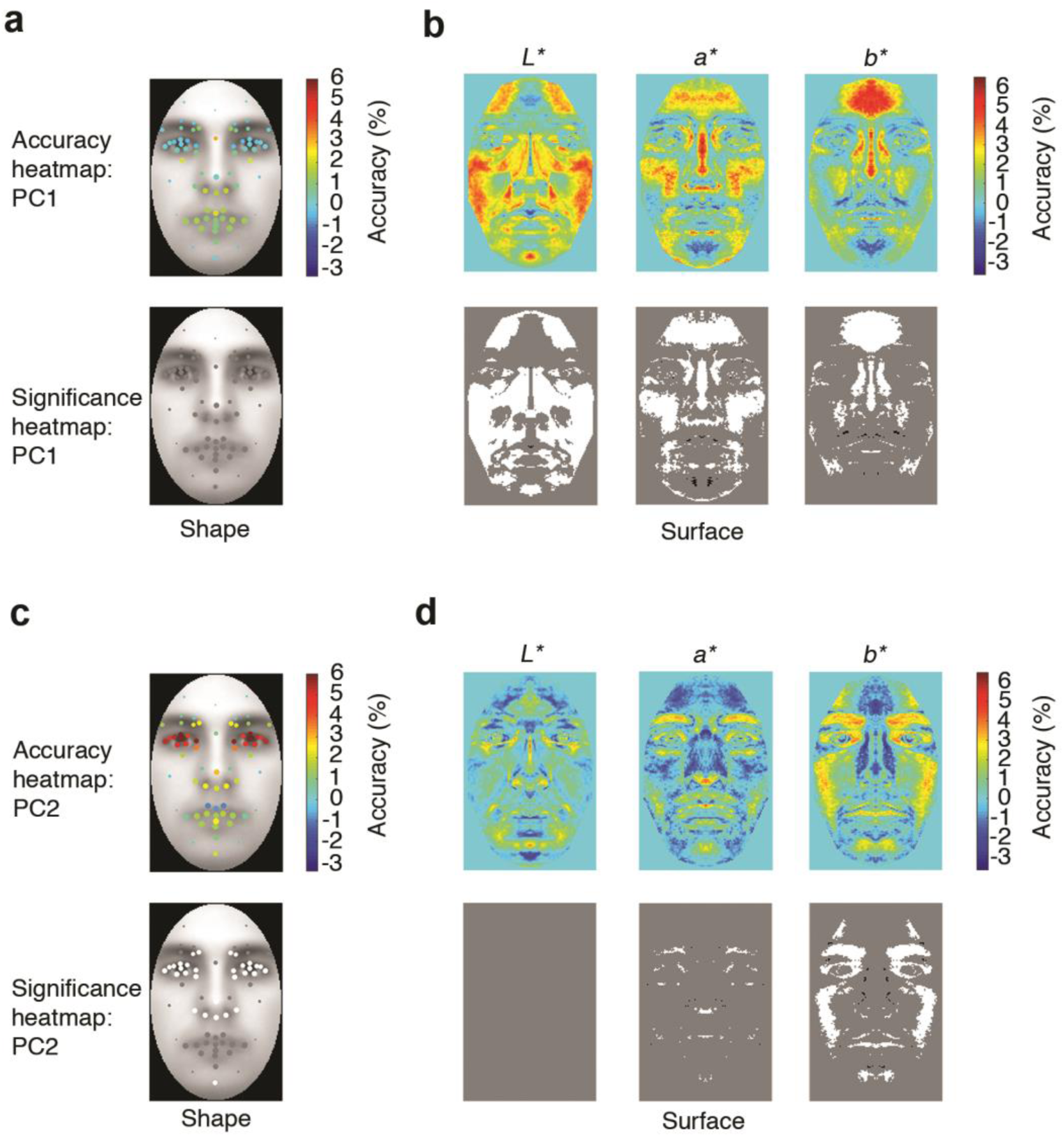
Heatmaps of differences in reconstruction accuracy for the first two principal components (PC1 and PC2) in the participant space. Heatmaps of such differences and their significance are constructed for (a, c) every fiducial point in shape, and (b, d) every pixel and color channel. In significance heatmaps, white/black indicates information more accurately retrieved in participants with positive/negative PC scores (FDR, *q* < .05) across fiducial points, for shape, and across pixels, for surface (gray marks the lack of significant differences). The relative size of fiducial points corresponds to their location variability across faces (*SD*). (The gray face images in (**a**) and (**c**) were artificially generated for illustration purposes in Matlab)

Thus, the present results confirm that aging plays an important role in face representation. However, visual discrimination ability, such as that captured by the FOJT, play a comparatively larger role in structuring face representations across participants. Moreover, such representations encode primarily eye shape and color information of faces, indicating again the importance of these features for face perception.

## Discussion

The present study sought to elucidate the impact of healthy aging on the visual representations underlying face processing by using a novel image reconstruction approach. Our investigation revealed the pictorial content associated with face perception in YAs and OAs as well as, critically, a variety of differences in shape and surface representation across age groups. More specifically, facial features that are crucial for facial identification, such as the shape of the eyes and the nose as well as the color of the forehead and the cheeks, were more accurately represented by YAs than OAs.

Our finding of decreased representational accuracy for shape in OAs compared to YAs, is consistent with previous studies arguing for an aging-related decline in the perception of face shape (Chaby et al., 2011; Murray et al., 2010; Slessor et al., 2013). Further, our results are convergent with recent studies pointing to higher reliance on visual information from the eye region in younger versus older adults (Chan, Chan, Lee, & Hsiao, 2018; Creighton, Bennett, & Sekuler, 2018). Regarding the role of surface information, we found that luminance properties were less accurately recovered for OAs compared to YAs, in agreement with poorer contrast sensitivity to face information in the former (Owsley & Sekuler, 1981). Importantly, our analyses provided novel evidence that aging impacts facial color representations, with the reconstruction of information from the red-green and yellow-blue channels being less accurate in the OA group, in particular around the forehead and cheek areas.

Thus, the present findings not only serve to confirm a number of hypotheses in the field but crucially, reveal novel aspects of visual representations undergoing aging-related changes. Specifically, by employing a data-driven approach that is not constrained by pre-defined hypotheses or by the use of artificially-manipulated features, our work provides unique insight into how aging impacts the pictorial content of face representations. Critically, the visual features uncovered here can account for the decline of face identification in older adults. For instance, the recognition of facial identity, gender, emotion and even animacy rely on visual information from the eye area (Balas & Horski, 2012; Nestor, Vettel, & Tarr, 2008; Schyns, Bonnar, & Gosselin, 2002; Sekuler, Gaspar, Gold, & Bennett, 2004; Smith, Cottrell, Gosselin, & Schyns, 2005) as well as on facial color (Benitez-Quiroz, Srinivasan, & Martinez, 2018; Dupuis-Roy, Fiset, Dufresne, Caplette, & Gosselin, 2014; Nestor, Plaut, & Behrmann, 2013). Thus, a decrease in the fidelity of corresponding representations is higly likely to impact multiple types of recognition including identification.

At the same time though, we found that age only accounted for a relatively small amount of data variance across participants. Instead, the largest components of variance were explained by individual differences captured by a different task, the different-view FOJT. The performance on this task, in which participants are asked to perceive faces from different viewpoints and to identify the odd one, has been suggested to reflect the ability to process feature conjunctions, which is necessary to discriminate between stimuli possessing overlapping features (Behrmann et al., 2016; Lee et al., 2005, 2006). Further, an evaluation of the data revealed extensive differences in both shape and surface perception across participants, independent of age. Thus, collectively, individual differences in visual representations appear to impact face perception to a greater extent than aging.

Regarding the nature of the information retrieved above, we note that surface information was systematically retrieved more accurately than shape information in all participants, irrespective of age. This finding is consistent with previous results demonstrating better sensitivity to surface than shape in human participants but not in a theoretical observer, which was capable of retrieving both types of information with equivalent levels of success (Nemrodov et al., 2019). More generally, these results are consistent with the dominant role of surface properties in face recognition (Burton, Jenkins, Hancock, & White, 2005; Kaufmann & Schweinberger, 2008; Russell, Sinha, Biederman, & Nederhouser, 2006; Vuong, Peissig, Harrison, & Tarr, 2005). As a caveat to this conclusion, we acknowledge that the present investigation did not consider three-dimensional face shape (Jiang, Blanz, & O’Toole, 2009) but only two-dimensional information. Hence, it is possible that, by facilitating the recovery of overall shape information, additional 3D cues would lead to more precise representations (i.e., at least as accurate as those of surface). Further, to better disentangle the contribution of shape and surface properties to face perception, future work may benefit from using artificial images that control the two types of information independently - but see evidence (Balas & Pacella, 2015) pointing to the disadvantage of artificial stimuli in the study of face recognition.

Another important avenue for future research concerns the role of the own-age bias in face processing, the finding that recognition memory for faces of one’s own age group is superior to memory for faces of another age (Bartlett & Leslie, 1986; Rhodes & Anastasi, 2012). Yet, the perceptual nature of this bias, though plausible, is still unclear (Wiese, Komes, & Schweinberger, 2013). Hence, the application of image reconstruction to faces of one own’s age and of different ages could shed light on the origin of this bias and also serve to disentangle its effect from that of a general decline in face recognition due to aging.

Last, from a methodological standpoint, it is important to highlight the current application of image reconstruction. Previous work, whether targeting faces (Chang et al., 2017; Lee & Kuhl, 2016; Nemrodov, Niemeier, Patel, & Nestor, 2018; Zhan et al., 2019) or other visual categories (Miyawaki et al., 2008; Naselaris, Prenger, Kay, Oliver, & Gallant, 2009; Nishimoto et al., 2011), has employed relatively small participant samples (e.g., 2 - 14) and has been restricted to neurologically healthy young adults (e.g., 20 – 30 years old), thereby not addressing the validity and the utility of image reconstruction across the wider population. Our work addresses this limitation by demonstrating that image reconstruction can be applied successfully to data collected from a cohort of 45 participants of varying age and perceptual ability. Further, it suggests that the current methodological framework may be successfully with other populations experiencing impaired visual recognition and visual distortions due to mental illness or brain damage.

In summary, the current work takes an important step toward elucidating the impact of healthy aging on the visual content underlying face perception. Our investigation reveals a wealth of visual information negatively impacted by aging and spanning both facial shape and surface properties. At the same time, our results put into perspective the impact of aging by demonstrating its limited extent when compared to that of overall individual variability. Last, the present work validates the applicability of image reconstruction to a broader population and demonstrates its utility in illuminating both group-level and individual differences in visual perception.

