## Supplemental Materials for "Image Reconstruction Reveals the Impact of Aging on Face Perception"

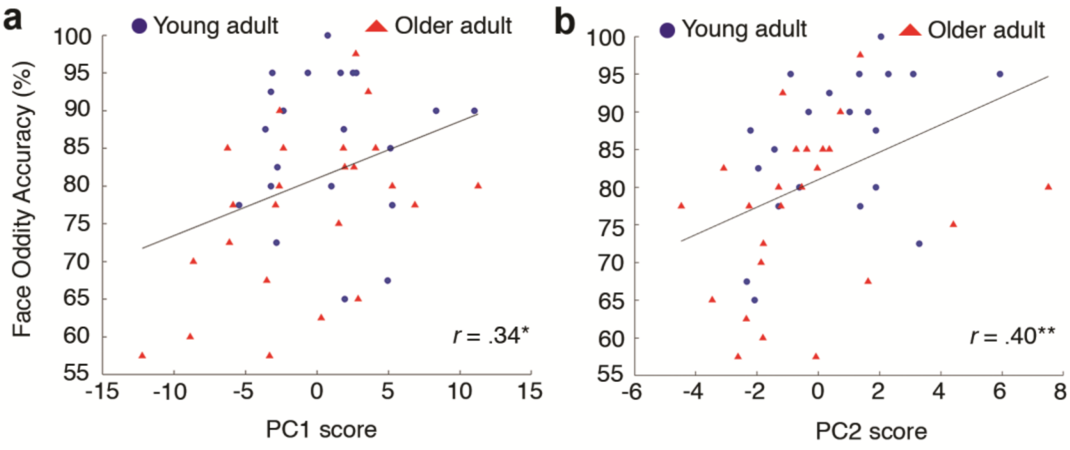


*Figure S1.* The relationship between different-view face oddity accuracy and principal component (PC) scores from (a) the first PC and (b) the second PC. The black line represents the fitted linear trend (* $p < .05$; ** $p < .01$).
